# Nonlinear hippocampal coding of the pair-bonded partner in prairie voles

**DOI:** 10.1101/2025.10.10.681589

**Authors:** Jiayan Liu, Kisa Watanabe, Miyuki Miyano, Mikuru Kudara, Eriko Kato-Ishikura, Sena Iijima, Kinjiro Takeda, Ryosuke Koike, Takeshi Nagahiro, Kotaro Yamashiro, Larry J. Young, Yuji Ikegaya, Nobuyoshi Matsumoto

## Abstract

Neural representation of specific opposite-sex partners remains largely unexplored. Using monogamous prairie voles, we demonstrated that partner preference required the ventral hippocampus, in which 3–4 Hz oscillations were enhanced following partner interaction. We identified ventral hippocampal neurons responding selectively to the pair-bonded partner and stranger, which encoded identity-related information in a nonlinear manner, shedding light upon how mammalian brains create distinct neural representations of romantic opposite-sex partners.

## Main

Prairie voles (*Microtus ochrogaster*) are socially monogamous rodents that form long-lasting pair bonds and exhibit strong partner preferences, making them a powerful model for scrutinizing selective social attachment ^1,2^. Previous studies have shown that this partner preference behavior involves multiple brain regions such as the nucleus accumbens (NAc) with its dopamine signaling playing a key role in bond formation and maintenance ^3,4^.

The NAc receives diverse inputs, including reward-related signals from the prefrontal cortex and contextual information from the hippocampus ^5,6^. In particular, the ventral hippocampus in mice and rats is essential for social memory, in which not only the familiarity of conspecifics but also the sex is encoded ^7–9^.

Here, we pharmacologically inactivated ventral hippocampal neural activity to address whether it is necessary for partner preference in male prairie voles. We then recorded ventral hippocampal neural activity during social interactions to identify neural correlates of partner-specific responses.

### Neural activity of the ventral hippocampus is required for partner preference behavior

To investigate partner preference in male prairie voles, we established an open field behavioral paradigm to assess partner preference in prairie voles. Transparent stimulus chambers containing partner and stranger females were positioned on opposite sides of the field, allowing the male subject to move freely. Males exhibited significantly longer interaction times with partner females compared to stranger females (Fig. 1a).

**Figure 1.**
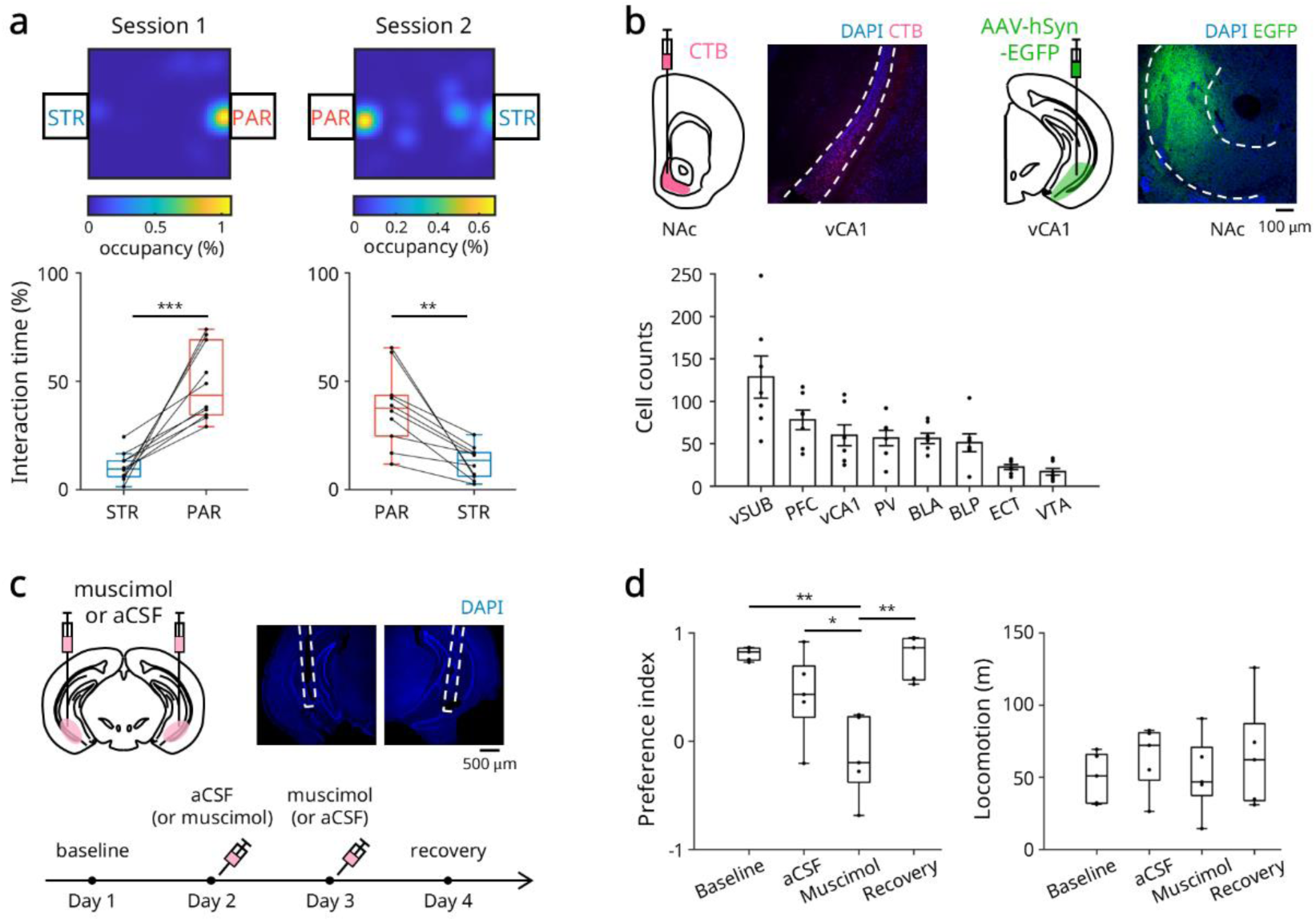
The ventral hippocampus is required for partner preference in male prairie voles. ***a***, Partner preference test. *Top*: Representative heatmaps depicting the proportion of time spent by male prairie voles at each location. Hot color indicates A partner female is located in a small chamber at one side, whereas a stranger female is in a chamber at the other side. The locations of the female voles are swapped from the Session 1 to 2 so that the possibility of place preference can be excluded. *Bottom*: The interaction time with the partner or stranger female, averaged across animals. Regardless of the female location, males exhibit a consistent preference for their partner (*n* = 10 prairie voles, \*\*\**P* < 0.001, paired *t*-test). ***b***, *Top*: (*Left*) Schematic of retrograde tracer (CTB) injection into the NAc and representative image showing labeled neurons in vHPC (*red*). (*Right*) Schematic of anterograde tracer (AAV-hSyn-EGFP) injection into vHPC and resulting projections to the NAc (*green*). *Bottom*: Quantification of retrogradely labeled cells in the NAc- upstream regions (*n* = 7 prairie voles). ***c***, Muscimol or aCSF was bilaterally injected into the ventral hippocampus of male prairie voles. Top: (*Left*) Illustration of the injection experiment. (*Right*) Representative DAPI-stained brain slice showing the injection site in the ventral hippocampus; the dashed white line indicates cannula placement. *Bottom*: Experimental timeline of the pharmacological manipulation. ***d***, *Left*: Preference index of male voles in each group. The preference index is defined as (partner interaction time – stranger interaction time) / (partner interaction time + stranger interaction time). The muscimol group shows a significantly reduced preference index compared to baseline, aCSF, and recovery groups (*n* = 5 prairie voles, \**P* < 0.05, \*\**P* < 0.01, paired *t*-test followed by Holm correction). *Right*: Locomotor activity of each group. No significant difference in locomotion across groups (*n* = 5 prairie voles, paired *t*-test followed by Holm correction). *Abbreviations*: STR, stranger; PAR, partner; CTB, cholera toxin subunit B; NAc, nucleus accumbens; vHPC, ventral hippocampus; aCSF, artificial cerebrospinal fluid.

Given that the NAc potentially mediates partner preference in prairie voles ^3,10^, we addressed how partner information is represented in the brain. As the NAc integrative function likely depends on afferent inputs conveying socially relevant information, we sought to identify upstream brain regions projecting to the NAc in prairie voles. To this end, we performed retrograde tracing by injecting cholera toxin B into the NAc of males and revealed eight major upstream regions, including the ventral CA1 (vCA1) and ventral subiculum (vSUB) of the hippocampus, with the ventral subiculum showing the highest density of labeled neurons (Fig. 1b, S1a); note that the ventral hippocampus has been implicated in processing social memory and contextual information in mice ^7,9^. We collectively refer to vCA1 and vSUB as the ventral hippocampus, hereafter. Anterograde tracing further confirmed direct projections from the ventral hippocampus to the NAc (Fig. 1b, S1b).

To test whether neural activity in the ventral hippocampus is required for partner preference, we conducted pharmacological inactivation experiments. Bilateral cannulas were implanted into the ventral hippocampus, and muscimol (*i.e.*, a GABA_A_ receptor agonist) or artificial cerebrospinal fluid (aCSF) as a control was injected on alternating days (Fig. 1c). Muscimol treatment significantly reduced the partner preference index relative to the baseline and aCSF control, with preference restored following recovery (Fig. 1d). These findings suggest that the ventral hippocampus is essential for partner preference in prairie voles.

### Cells responsive to partners or strangers in the ventral hippocampus

To further scrutinize how the vole ventral hippocampal neurons responded during interactions with a partner or a stranger, we conducted electrophysiological recordings using silicon probes. Silicon probes were unilaterally implanted into the ventral hippocampus in male prairie voles. After the surgery and recovery, the male prairie voles were allowed to explore the open field with nose-poke sensors attached to stimulus females to precisely define interaction onset (Fig. 2a). We identified functionally distinct neuronal populations: partner (PAR) onset ON cells that increased firing during partner-associated nose-pokes, stranger (STR) onset ON cells responsive to stranger-interactions, and Both onset ON cells responsive to both females (Fig. 2b, c). We also found corresponding OFF cells that decreased firing during interactions (Fig. 2b, d), with similar patterns for interaction offset (Fig. S2). These findings indicate that the ventral hippocampus exhibits temporally specific neural representations of social interactions with identified females.

**Figure 2.**
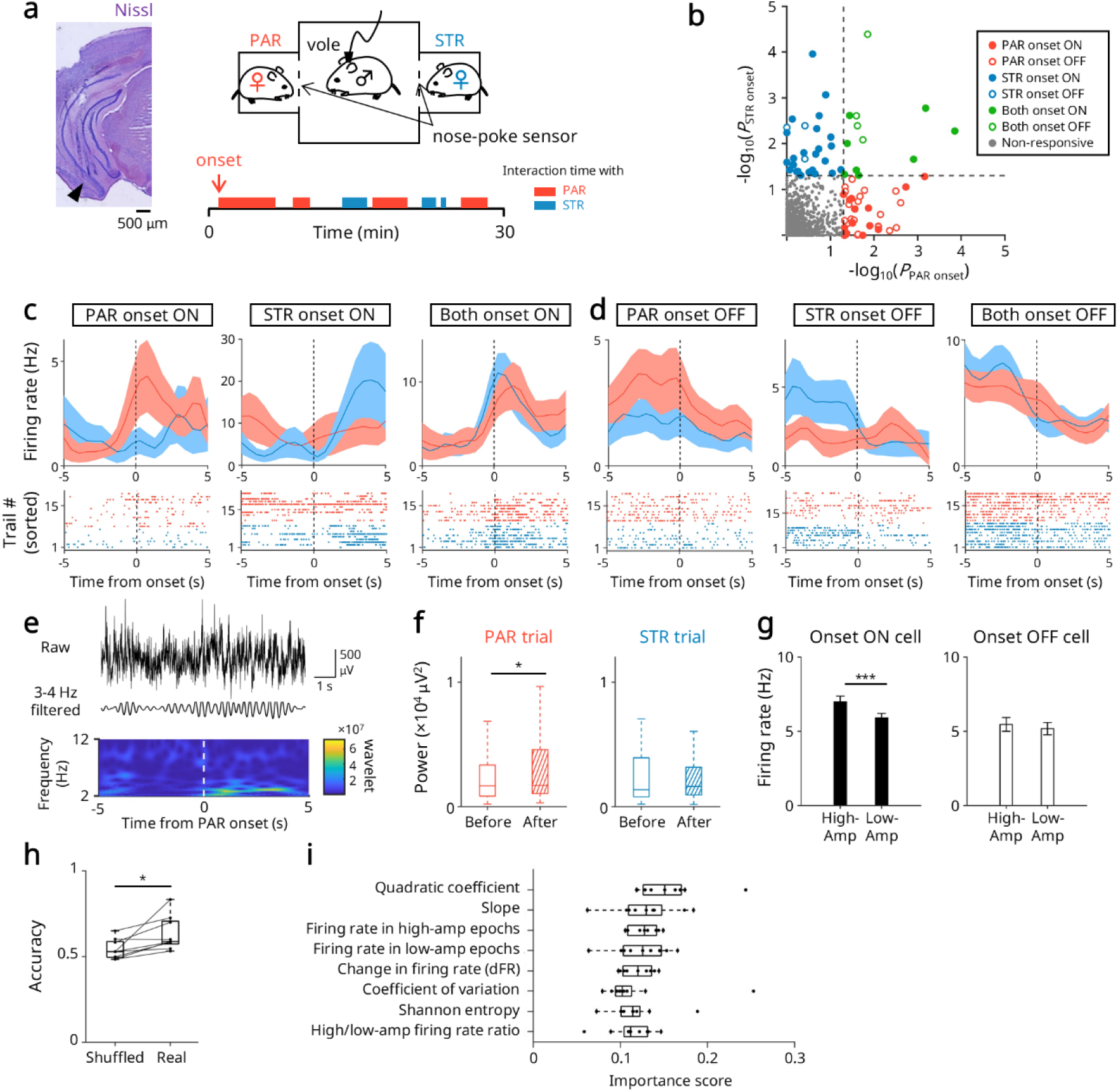
Neural correlates of partner and stranger females in the ventral hippocampus in male prairie voles. ***a***, *Left*: A silicon probe is unilaterally implanted into the ventral hippocampus of male prairie voles. The black arrow indicates the probe track. *Right*: Timeline of electrophysiological recording. Nose-poke sensors are attached to the chambers containing each female. The time at which the male began nose-poking toward the partner (*red*) or stranger (*blue*) was defined as the partner or stranger onset. ***b***, For each recorded cell, the significance (*P* value) of the difference in firing rates before and after the partner onset is transformed as –log_10_(*P*) and plotted on the x-axis. The same computation was performed for the stranger onset and plotted on the y-axis (*n* = 1078 cells from 5 prairie voles, Wilcoxon rank-sum test). ***c***, Some neurons increase their firing rates specifically at the partner onset, stranger onset, or both. These cells are named PAR (partner) onset ON, STR (stranger) onset ON, and Both onset ON cells, respectively. ***d***, Some other neurons decrease their firing rates specifically at the partner onset, stranger onset, or both. These are named PAR onset OFF, STR onset OFF, and Both onset OFF cells, respectively. ***e***, *Top*, A representative raw trace of local field potentials (LFPs) 5 seconds before and after partner onset. *Middle*, A representative filtered (3–4 Hz) trace that corresponds to the *top* trace. *Bottom*, A scalogram based on the wavelet transformation of the LFPs 5 s before and after the partner onset, showing enhanced 3–4 Hz oscillations following the onset. ***f***, Quantification of 3–4 Hz power 5 s before *vs*. after the partner onsets, confirming a significant increase in 3–4 Hz power following the partner onset, while no such change occurs for the stranger onset (*n* = 118 partner onsets, *n* = 101 stranger onsets, \**P* < 0.05, paired *t*-test). ***g***, Across all onset ON cells, the firing rates during high-amplitude epochs are significantly higher than those during low-amplitude epochs (*n* = 557 firing events from 2–16 trials in 2–12 onset ON cells after the partner onsets, *n* = 307 firing events from 2–16 trials in 1–14 onset OFF cells after the partner onsets, \*\*\**P* < 0.001, Student’s *t*-test). ***h***, Interaction partners are predicted using firing patterns during the 5-s period before the onset. The prediction accuracy using real labels is significantly higher than that with shuffled labels (*n* = 9 recordings from 5 prairie voles, \**P* < 0.05, paired *t*-test). ***i***. Feature importance for decoding interaction partners (*n* = 9 recordings from 5 prairie voles). *Abbreviations*: PAR, partner; STR, stranger.

In line with previous work demonstrating that ventral hippocampal neurons in mice showed selective activity near familiar or novel individuals and responded differentially to individuals of different sexes ^7,8^, we identified cells that transiently altered firing at moments of social approach or departure. The presence of neurons that respond oppositely to the same individual may enhance the contrast of partner-specific coding. Notably, approach-and departure-responsive neurons have also been reported in the NAc, where approach cells expand in number with pair bond formation ^10^. Given that there are projections from the ventral hippocampus to the NAc, these regions may act in concert to support social preference behavior.

### Enhanced 3–4 Hz oscillations following partner interaction onset

Beyond single-unit activity, we scrutinized broader network dynamics in the ventral hippocampus by analyzing local field potentials (LFPs). LFP analysis revealed a significant enhancement of 3–4 Hz oscillations, a novel frequency band, in the ventral hippocampus following partner onset, without such increase after stranger onset (Fig. 2e). The power in other frequency bands did not change after partner onset, although theta (6– 12 Hz) and gamma (50–80 Hz) bands were enhanced following stranger onset (Fig. S3). When 3–4 Hz oscillatory amplitudes were used to segment the post-onset period into high- and low-amplitude epochs relative to the mean, onset ON cells exhibited significantly higher firing rates during high-amplitude epochs than low-amplitude epochs. In contrast, onset OFF cells did not exhibit such significant differences, suggesting that enhanced 3–4 Hz oscillations temporally gate partner-specific activation of ON cells (Fig. 2f, S4, S5).

Enhanced 3–4 Hz oscillations following partner interaction are indicative of encoding social familiarity. This frequency band overlaps with eta or high-delta oscillations observed in rodent hippocampus ^11^ and may be generated via local inhibitory-excitatory interactions or long-range prefrontal-thalamic inputs ^12^. Notably, similar 4 Hz oscillations have been implicated in social memory via prefrontal-ventral tegmental and hippocampal phase coupling ^13^. In contrast, stranger onset increased theta (6–12 Hz) and high-gamma (50–80 Hz) activity—oscillations associated with exploratory encoding and novelty detection ^14,15^. These results suggest that partner- and stranger-directed behaviors engage distinct hippocampal network dynamics, with 3–4 Hz rhythms selectively coordinating recognition processes for social familiarity and ON cell activity, leading to the partner representation in prairie voles.

### Decoding partners or strangers from pre-onset firing patterns

While post-onset neural activity in the hippocampus characterized opposite-sex partners, we questioned whether pre-onset firing patterns of onset ON cells encoded information predictive of upcoming interaction partners. To this end, we applied machine learning models combining Random Forest and XGBoost to extract features from the firing activity in the 5-second period preceding the social onset. The decoding accuracy for real female labels (*i.e.*, a partner or a stranger) was significantly higher than that of shuffled labels, indicating that pre-onset neural activity reliably predicted the identity of the interaction partner (Fig. 2g).

We then examined what feature in the pre-onset neural activity in the male vole hippocampus was informative for decoding female identity. The features input to the model were calculated based on the firing rates of each onset ON cell during the 5-second period preceding the onset. When these features were ranked by importance, the quadratic coefficient and slope of firing rate emerged as the top two features (Fig. 2h). Additionally, the firing rate in high-amplitude 3–4 Hz epochs and firing rate in low-amplitude 3–4 Hz epochs (discussed above) also constituted important features. These findings suggest that temporal changes in firing rate and 3–4 Hz-related firing patterns contain predictive information of female identity for interaction.

Nonlinear and temporal derivative features of neural activity predict the future sexual counterpart. Our findings align with recent theoretical and empirical work showing that nonlinear transformations of neural responses, particularly quadratic components, can encode behaviorally relevant information inaccessible to linear decoders ^16^. In parallel, slope-sensitive burst firing—previously observed in both biophysical neuron models ^17^ and point neuron simulations ^18^—likely underlies the strong predictive power of the slope feature. These features may reflect distinct encoding strategies: the quadratic term captures curvature or non-monotonic dynamics, while the slope reflects the temporal directionality of firing rate changes preceding behavior. Our findings extend these prior observations to a novel social context, where pre-onset firing predicts social engagement targets. Notably, features like entropy and coefficient of variance, known to reflect variability and coding capacity ^19,20^, also showed moderate importance, suggesting a potential role in modulating encoding precision. Altogether, these results support the utility of nonlinear and temporal derivatives in decoding upcoming social behavior from neural activity.

Collectively, our findings bring about insights into how mammalian brains create distinct neural representations of romantic opposite-sex individuals at the single-cell and population levels. Deciphering how the brain encodes and maintains affiliative relationships with close social partners will open new avenues for neural underpinnings of sexuality beyond sociality.

## Author contributions

J.L., Y.I. and N.M. conceived the project and designed the study; J.L., M.M. and N.M. performed the experiments; J.L., K.W., M.K., E.K-I., S.I., K.T., R.K., T.N., K.Y. and N.M. analyzed the data; J.L, Y.I. and N.M. oversaw and managed the project; J.L., Y.I., and N.M. wrote and sophisticated the manuscript. All approved the final version of the manuscript.

## Materials and Methods

### Ethics

All experiments were performed with the approval of the animal experimental ethics committee at the University of Tokyo (approval number: P3-1) and in accordance with the NIH guidelines for the care and use of animals.

### Animals

Prairie voles were gifted from Dr. Larry J. Young at Emory University and Dr. Shinichi Mitsui at Gunma University, and bred in our laboratory. Animals aged 5 weeks or older were used for experiments. They were group-housed under controlled temperature and humidity conditions (22 ± 1°C, 55 ± 5%) with a 12-h light/dark cycle (lights on from 7 a.m. to 7 p.m.) and given free access to food and water. Male prairie voles were used as subjects in all behavioral and electrophysiological experiments, while female prairie voles were presented to males as social stimuli in the partner preference test.

### Behavioral assay

An apparatus used in this study was a cardboard field (40 cm (in length) × 40 cm (in width) × 25 cm (in height)) with transparent acrylic chambers (8 cm (L) × 8 cm (W) × 15 cm (H)) placed on both sides. The field was covered with bedding materials. One side of each acrylic chamber facing the field had 36 holes (4 columns × 9 rows, 1 cm diameter) for nose-poke contacts. The behavioral test consisted of two sessions. In Session 1, the partner female was placed in either the left or right acrylic chamber, while a stranger female was placed in the opposite chamber. The male was allowed to freely move in the field and nose-poke for interaction with females. In Session 2, the positions of the partner and stranger females were switched, and the same experiment was conducted. Each session lasted 30-60 min. The field was surrounded by black curtains to maintain a dim environment. The male’s behavior was recorded at 60 fps using an infrared camera (Autocastle, Shenzhen Ailipu Technology, China).

Video recordings were first converted to 5 fps offline using ffmpeg (https://www.ffmpeg.org/), then processed frame-by-frame using Python. Interaction was defined as the male orienting toward and making contact with a female’s chamber. The number of frames showing interaction with either partner or stranger female was counted. The percentage of total frames spent in interaction with partner *vs*. stranger females was calculated and compared using the paired *t*-test (Fig. 1a). A preference index was calculated by dividing the difference between partner and stranger female interaction frames by their sum (Fig. 1b).

For pharmacological inhibition experiments, the male’s movement trajectory for each day was tracked using ImageJ. Locomotion was calculated from the change in male coordinates between frames.

### Injection

General anesthesia was induced with 4% (v/v in air) isoflurane and maintained with 1– 2% isoflurane. Veterinary ointment was applied to the eyes to prevent drying. The skin was sterilized with povidone-iodine and 70% ethanol before incision. After anesthesia, the prairie vole was mounted in a stereotaxic frame using ear bars and a nose clamp. The scalp was incised with surgical scissors, and holes (∼0.5 mm in diameter) were drilled using a high-speed dental drill (SD-102, Narishige, Japan).

The following materials were injected at 100 nL/min until a total volume of 200 nL using a Hamilton syringe: Alexa Fluor 555-conjugated cholera toxin subunit B (CTB; 1.0 mg/mL, C34776, ThermoFisher Scientific, USA) into the nucleus accumbens (2.2 mm anterior and 0.70 mm lateral to the bregma, and 4.5 mm beneath the brain surface) and adeno-associated virus (AAVdj-hSyn-EGFP; 1.4 × 10¹³ vg/mL) into the ventral hippocampus (3.3 mm posterior and 3.5–3.8 mm lateral to the bragma, and 3.5–4.3 mm beneath the brain surface). The scalp was sutured using surgical needles to protect the cranial window. Animals were sacrificed at least a week or two weeks after the surgery for CTB or AAV injection, respectively,

### Pharmacological inactivation

Under isoflurane anesthesia, the prairie vole’s scalp was incised and 0.8 mm diameter holes were drilled at coordinates of 3.3 mm posterior and 3.5–3.8 mm bilateral to the bregma using a high-speed drill. The dura was surgically removed. Two guide cannulae were inserted into the holes to a depth of 3.5–4.3 mm beneath the brain surface, and dummy cannulae were inserted into the guide cannulae for occlusion. The assembly was secured with dental resin (Provinice, Shofu, Japan) and cement (Super-Bond, Sun Medical, Japan).

For drug administration, the dummy cannulae were removed and injection cannulae were inserted through the guide cannulae. Either muscimol solution (0.1 mg/mL in artificial cerebrospinal fluid (aCSF)) or aCSF was infused into the ventral hippocampus bilaterally at 100 nL/min until a total volume of 200–300 nL. At least 1 min after the injection, the injection cannulae were removed and dummy cannulae were reinserted. Behavioral tests were begun 15 min after the cannula replacement.

On Day 1, behavioral test was conducted with the same protocol in the behavioral assay without any drugs administered to voles, which was regarded as a baseline experiment. On Day 2, either muscimol solution or aCSF was administered before the behavioral test. On Day 3, the alternative solution was administered before the test. On Day 4, the same behavioral test as on Day 1 was conducted to confirm recovery of partner preference.

### *In vivo* electrophysiology

128-channel silicon probes (P64-1-D (6 mm), Diagnostic Biochips, USA) were used for electrophysiological recordings. A custom-made microdrive to adjust the height of the silicon probe was manufactured using a 3D printer (MAX X43, Asiga, Australia).

Under isoflurane anesthesia, cranial windows (1.5 mm in diameter) were made at coordinates of 3.3 mm posterior and 3.5–3.8 mm bilateral to the bregma, after which the dura was surgically removed. The silicon probe tip was lowered 3.0–4.0 mm from the brain surface, and the space between the probe and brain surface was protected with biocompatible silicone adhesive (Kwik-Sil, World Precision Instruments, USA). The microdrive was then secured to the skull using dental cement and super bond.

Two stainless steel screw electrodes were implanted on the cerebellar surface as ground/reference electrodes^21^ and soldered to the ground/reference wires extending from the silicon probe. To prevent damage to the microdrive, a protective hood made of copper mesh was secured to the skull using adhesive and cement and further connected to the reference wire by soldering.

The silicon probe implanted in the prairie vole was connected to a data acquisition system (RHD recording system, Intan Technologies, USA) through its analog-to-digital converter. Neural signals from each channel were monitored in real-time using data acquisition software (RHX data acquisition software, Intan Technologies). The probe tip could be chronically lowered to the target region by turning the microdrive screw. To record ventral hippocampal neural activity, the silicon probe was lowered 35–280 μm until reaching the pyramidal cell layer of the ventral hippocampal CA1 area, with the aid of visual inspection of the presence of ripples (120–240 Hz). Recordings were conducted over several days.

The same open field and acrylic chambers as described in the behavioral assay were used in the behavioral electrophysiological experiment. Nose-poke sensors (PS-RL, O’hara, Japan) were attached to the acrylic chambers to precisely detect the time when the male vole interacted with females. When the male nose-poked, an infrared sensor was triggered, sending an analog signal to the data acquisition system through a poking sensor controller (PS-2020, O’hara). Additionally, the sensor identifier (*i.e.*, partner female side or stranger female side) was sent as serial output through Arduino Uno.

The behavioral task consisted of the Session 1 (10–30 min) with the partner female in one chamber and stranger female in the other, followed by the Session 2 (10– 30 min) with the females’ positions swapped. Neural activity was also recorded during pre-rest (30–60 min) and post-rest (30–60 min) periods, in which the recorded animal was in its home cage before and after the task, respectively. Throughout all recording periods, the animal’s behavior was filmed using a camera (MCM-303NIR-880-LED, Gazo, Japan) on the ceiling above the field. The camera’s strobe signals at a rate of 60 Hz were transmitted as an analog signal to the data acquisition system. Signals from the silicon probes, nose-poke sensors, and camera were recorded at a sampling rate of 30 kHz.

### Histology

Prairie voles underwent (1) CTB/virus injection experiments, (2) pharmacological inhibition experiments, or (3) electrophysiological experiments. Animals were perfused at the following timepoints: (1) 7 or 14 days after surgery for CTB or AAVdj-hSyn-EGFP injection, respectively, and (2, 3) immediately after completion of experiments. Animals were deeply anesthetized with isoflurane and perfused transcardially. After confirming complete anesthesia by absence of tail and toe pinch reflexes, ice-cold 0.01M phosphate-buffered saline (PBS) was perfused followed by 4% paraformaldehyde (in PBS). The brain was carefully removed and post-fixed overnight in 4% paraformaldehyde, then transferred to 30% sucrose solution. Subsequent procedure was below.

1. Brains were frozen on dry ice and sectioned at 50 μm using a cryostat (HM525NX, epredia, USA). Sections were stored in PBS, incubated in fluorescent DAPI solution (1:1000 in PBS; D1306, ThermoFisher) at room temperature for 90 min, washed in PBS for 10 min, and coverslipped using CC/Mount (Diagnostic BioSystems, USA).
2. Brains were frozen on dry ice and sectioned at 100 μm using a cryostat. Sections were stored in PBS, incubated in DAPI solution (1:1000 in PBS) at room temperature for 90 min, washed in PBS for 10 min, and coverslipped using CC Mount.
3. Brains were frozen on dry ice and sectioned at 50 μm using a cryostat. Sections were stored in PBS, mounted on slides using mounting solution (0.2% gelatin in PBS), and dried overnight. Cresyl violet staining was then performed. Mounted sections were hydrated in distilled water, then dehydrated through an ethanol series (50, 60, 70, 80, 90, 100%). Sections were defatted in PathoClean (161-28321, FUJIFILM Wako, Japan) for 2 min, then rehydrated through a reverse ethanol series (100, 90, 80, 70, 60, 50%) for at least 10 s each. After rinsing in distilled water, sections were stained in cresyl violet solution for 10 min in the dark. Once adequately stained, sections were differentiated in 95% ethyl acetate solution and 100% ethanol. After a final PathoClean step, sections were coverslipped using PathoMount mounting medium (164-28492, FUJIFILM Wako).

### Image acquisition and analysis

For experiments (1) and (2) (see above), images were captured using a confocal microscope (A1 HD25, Nikon, Japan) with 4×, 10×, and 20× objectives. The following parameters were used for Z-stack imaging. (i) 4×: 11.625 μm for the step size, 3 steps; (ii) 10×: 2.075 μm for the step size, 5 steps; (iii) 20×: 0.6 μm for the step size, 5 steps. Z-stack images were averaged and contrast-adjusted using ImageJ (https://imagej.net/software/fiji/).

For experiment (3) (see above), brightfield images were captured using a light microscope (BZ-X800, Keyence, Japan) with a 2× objective. These images were used to identify the track of each recording electrode.

Images captured with the 20× objective were Z-stack-averaged and contrast-adjusted using ImageJ. Brain regions were manually outlined based on DAPI staining, and areas of the regions of interest (ROIs) were calculated. CTB-positive cells within the outlined regions were manually counted and divided by the ROI area to calculate cell density.

### Data analysis for electrophysiology

For electrophysiological data (digitized at a sampling rate of 30 kHz), spike sorting was first performed automatically using Kilosort2 (https://github.com/jamesjun/Kilosort2) in MATLAB 2021a (MathWorks, USA). Clusters were manually adjusted using Phy (https://github.com/cortex-lab/phy); putative neuronal units were isolated based on autocorrelation and cross-correlation of spike times.

Nose-poke time was detected based on analog signals sent by sensors attached to partner and stranger female cages. Due to continuous nose-poking by the male prairie vole, nose-pokes separated by more than 30 seconds were considered individual trials. The first poke time in a trial was defined as the onset, while the last poke time was defined as the offset.

For each cell, spike time was collected for 5 s before and after partner and stranger female onsets. Average firing rates were calculated for the 5-s periods before and after the onset. After cells that did not fire during the 10-s window around the onset were excluded from subsequent analysis, average firing rates during pre-onset and post-onset periods across all partner or stranger trials were compared using the Wilcoxon rank-sum test. Cells were classified as follows:

1. PAR onset ON cells: firing was significantly increased only after the partner onset.
2. PAR onset OFF cells: firing was significantly decreased only after the partner onset.
3. STR onset ON cells: firing was significantly increased only after the stranger onset.
4. STR onset OFF cells: firing was significantly decreased only after the stranger onset.
5. Both onset ON cells: firing was significantly increased after the partner and stranger onsets.
6. Both onset OFF cells: firing was significantly decreased after the partner and stranger onsets.
7. Non-responsive cells: firing was not significantly changed after partner or stranger onset.

Similar analysis was performed for offset responses. Cells were classified as PAR offset ON/OFF cells, STR offset ON/OFF cells, or Both offset ON/OFF cells based on significant changes in firing rate after the offset.

Single-channel LFP signals were downsampled to 500 Hz for further analysis. The fast Fourier transform was applied to LFP segments from 5 s before and after partner or stranger female onsets to estimate the power spectral density in a frequency-based domain. The power was calculated as the area under the curve for each frequency band (3–4 Hz (sub-theta), 6–12 Hz (theta), 20–30 Hz (beta), 30–50 Hz (low gamma), 50-80 Hz (high gamma)). Changes in the power before and after the onset were evaluated using paired *t*-tests.

Amplitude was calculated using the Hilbert transform for LFP signals 5 s before and after partner and stranger female onsets. For all partner onsets in a recording session, average amplitude was calculated for the 5-s periods before or after onset. The sub-periods with amplitude greater than the average amplitude were defined as high-amplitude epochs, while those below the average amplitude were named low-amplitude epochs. Similar calculations were performed for stranger female onsets. High-amplitude and low-amplitude epochs were defined for four 5-s periods (*i.e.*, before and after the partner onset, and before and after the stranger onset), and firing rates for each cell type were calculated.

Decoding of females (*i.e.*, the partner or stranger) was performed using the firing activity of all ON cells recorded in a single session. First, the spike times of ON cells during the 5-second period preceding partner and stranger interaction onsets were collected. The firing rate was then calculated in 1-second bins. Next, features were extracted from the firing activity of a single cell during a single trial and used as input parameters for a machine learning model, with the onset type (*i.e.*, partner or stranger) serving as the true label. We divided the 5-second period preceding the onset into 1-second bins and calculated the firing rate for each bin. The firing rate (FR) from *k* to *k`−`1* seconds before the onset was defined as FR*_k_* (where *k* = 1, 2, 3, 4, 5). The following eight features were used for prediction: (1) the change in firing rate (dFR), calculated as dFR = FR_1_ - (FR_2_ + FR_3_ + FR_4_ + FR_5_)/4, (2) the slope obtained from a linear regression of FR*_k_* against time, (3) the quadratic coefficient from a quadratic approximation of FR*_k_* against time, (4) the coefficient of variation (CV) of FR*_k_*, (5) the Shannon entropy of FR*_k_*, (6) the firing rate during high-amplitude epochs (for 3–4 Hz oscillations) within the 5-second period before onset, (7) the firing rate during low-amplitude epochs (for 3–4 Hz oscillations) within the 5-second period before onset, (8) the high-amplitude and low-amplitude firing rate ratio calculated as (firing rate in high-amplitude epochs – firing rate in low-amplitude epochs) / (firing rate in high-amplitude epochs + firing rate in low-amplitude epochs). Classification was performed using RandomForestClassifier and XGBClassifier from the Python sklearn library. Predictions were combined using soft voting with weighted averaging in a 1:1 ratio to yield the final results. Decoding was evaluated using classification accuracy, which was computed via 10-fold cross-validation.

## Acknowledgments

The authors sincerely thank Shota Morikawa, Haruki Takeuchi, Ayako Ishigaki, Kano Izumo, Kazuki Kakehi, and Haruka Suzuki for their technical assistance and insightful comments. This work was supported by JST ERATO (JPMJER1801), the Institute for AI and Beyond of the University of Tokyo, JSPS KAKENHI (22K21353), AMED Brain/MINDS 2.0 (24wm0625207s0101; 24wm0625401h0001; 24wm0625502s0501) (to Y. Ikegaya), JSPS KAKENHI (25K18705) (to N. Matsumoto), JSPS KAKENHI (22KJ1134) (to J. Liu).

## References

1. Carter, C. S., Witt, D. M., Thompson, E. G. & Carlstead, K. Effects of hormonal, sexual, and social history on mating and pair bonding in prairie voles. Physiol. Behav. 44, 691–697 (1988).

2. Aragona, B. J. & Wang, Z. The prairie vole (Microtus ochrogaster): an animal model for behavioral neuroendocrine research on pair bonding. ILAR J. 45, 35–45 (2004).

3. Aragona, B. J., Liu, Y., Curtis, J. T., Stephan, F. K. & Wang, Z. A critical role for nucleus accumbens dopamine in partner-preference formation in male prairie voles. J. Neurosci. 23, 3483–3490 (2003).

4. Pierce, A. F. et al. Nucleus accumbens dopamine release reflects the selective nature of pair bonds. Curr. Biol. 34, 519–530.e5 (2024).

5. Ito, R., Robbins, T. W., Pennartz, C. M. & Everitt, B. J. Functional interaction between the hippocampus and nucleus accumbens shell is necessary for the acquisition of appetitive spatial context conditioning. J. Neurosci. 28, 6950–6959 (2008).

6. Sosa, M., Joo, H. R. & Frank, L. M. Dorsal and ventral hippocampal sharp-wave ripples activate distinct nucleus accumbens networks. Neuron 105, 725–741.e8 (2020).

7. Okuyama, T., Kitamura, T., Roy, D. S., Itohara, S. & Tonegawa, S. Ventral CA1 neurons store social memory. Science 353, 1536–1541 (2016).

8. Watarai, A., Tao, K. & Okuyama, T. Representation of sex-specific social memory in ventral CA1 neurons. Science 389, eadp3814 (2025).

9. Meira, T. et al. A hippocampal circuit linking dorsal CA2 to ventral CA1 critical for social memory dynamics. Nat. Commun. 9, 4163 (2018).

10. Scribner, J. L. et al. A neuronal signature for monogamous reunion. Proc. Natl. Acad. Sci. U. S. A. 117, 11076–11084 (2020).

11. Safaryan, K. & Mehta, M. R. Enhanced hippocampal theta rhythmicity and emergence of eta oscillation in virtual reality. Nat. Neurosci. 24, 1065–1070 (2021).

12. Roy, A., Svensson, F. P., Mazeh, A. & Kocsis, B. Prefrontal-hippocampal coupling by theta rhythm and by 2-5 Hz oscillation in the delta band: The role of the nucleus reuniens of the thalamus. Brain Struct. Funct. 222, 2819–2830 (2017).

13. Fujisawa, S. & Buzsáki, G. A 4 Hz oscillation adaptively synchronizes prefrontal, VTA, and hippocampal activities. Neuron 72, 153–165 (2011).

14. Buzsáki, G. Theta oscillations in the hippocampus. Neuron 33, 325–340 (2002).

15. Neves, L. et al. Theta and gamma oscillations in the rat hippocampus support the discrimination of object displacement in a recognition memory task. Front. Behav. Neurosci. 16, 970083 (2022).

16. Yang, Q., Walker, E., Cotton, R. J., Tolias, A. S. & Pitkow, X. Revealing nonlinear neural decoding by analyzing choices. Nat. Commun. 12, 6557 (2021).

17. Kepecs, A., Wang, X.-J. & Lisman, J. Bursting neurons signal input slope. J. Neurosci. 22, 9053–9062 (2002).

18. Miko, R., Scheunemann, M. M., Steuber, V. & Schmuker, M. Decoding the amplitude and slope of continuous signals into spikes with a spiking point neuron model. bioRxiv (2024) doi:10.1101/2024.05.20.594931.

19. Luczak, A. Entropy of neuronal spike patterns. Entropy (Basel) 26, (2024).

20. Akella, S. et al. Deciphering neuronal variability across states reveals dynamic sensory encoding. Nat. Commun. 16, 1768 (2025).

21. Takeda, K. et al. Ramelteon coordinates theta and gamma oscillations in the hippocampus for novel object recognition memory in mice. J. Pharmacol. Sci. 158, 121–130 (2025).

