## Supplementary figures and images for "Nonlinear hippocampal coding of the pair-bonded partner in prairie voles"

### Figure S1

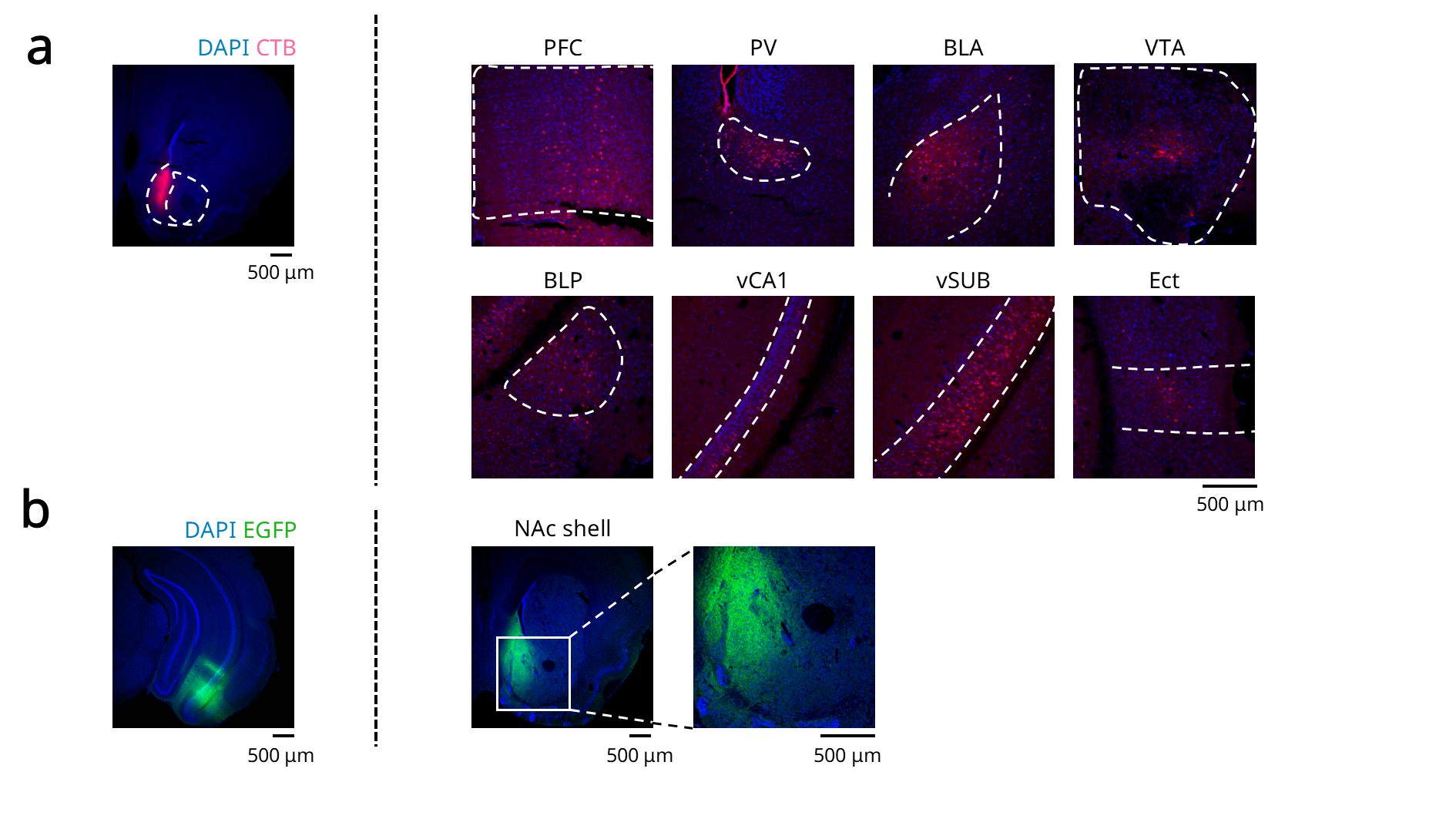

### Figure S2

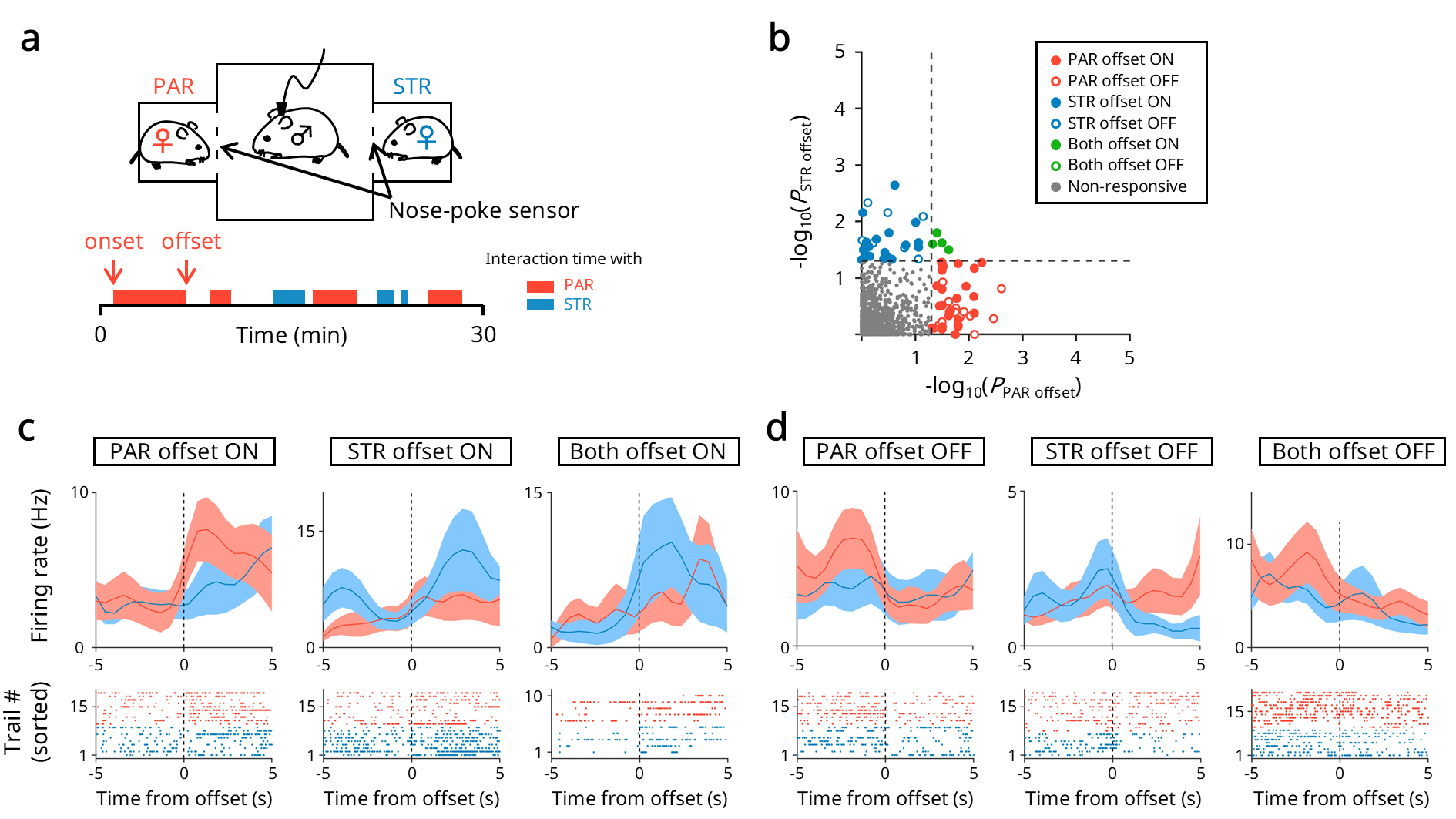

### Figure S3

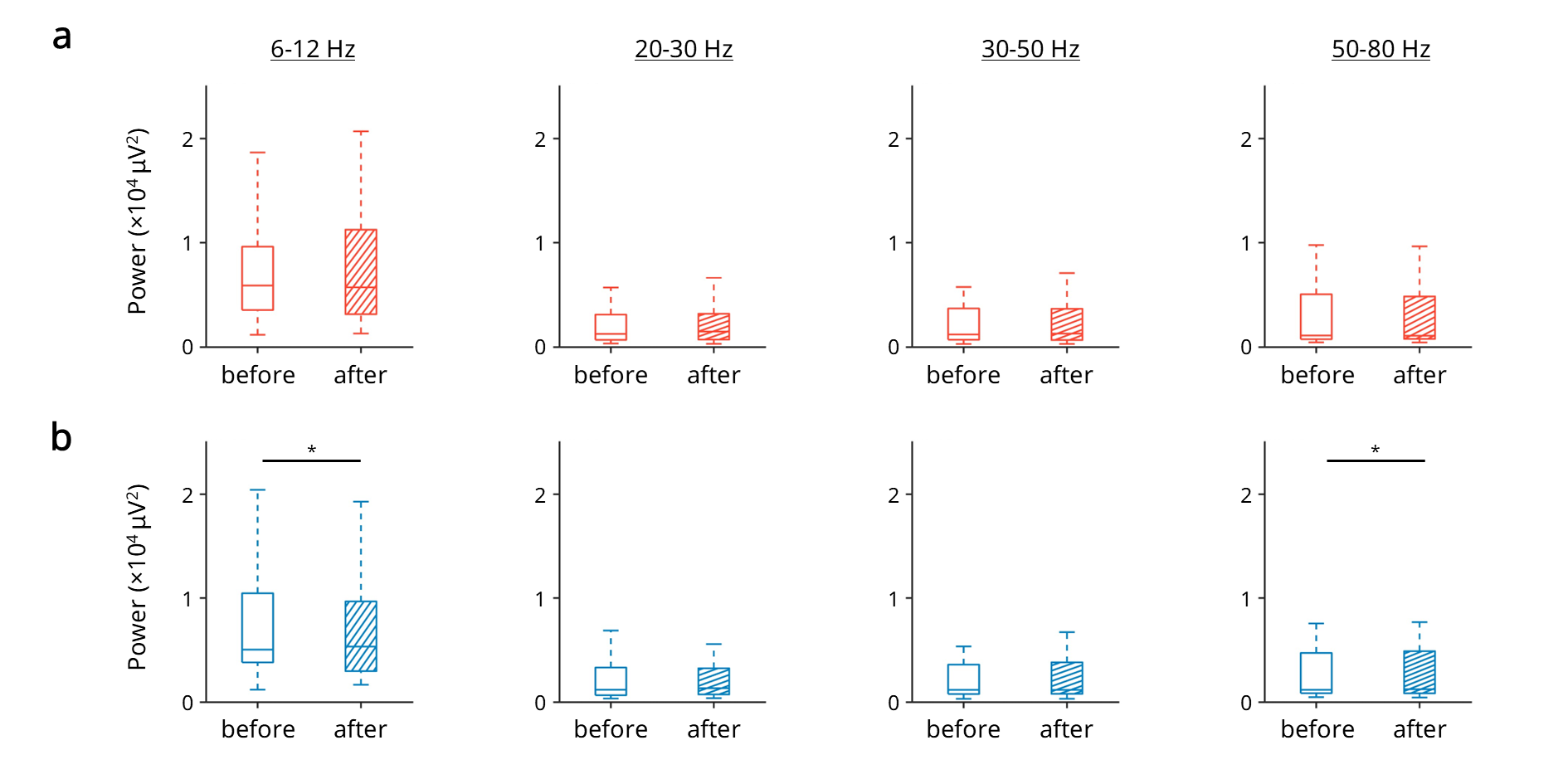

### Figure S4

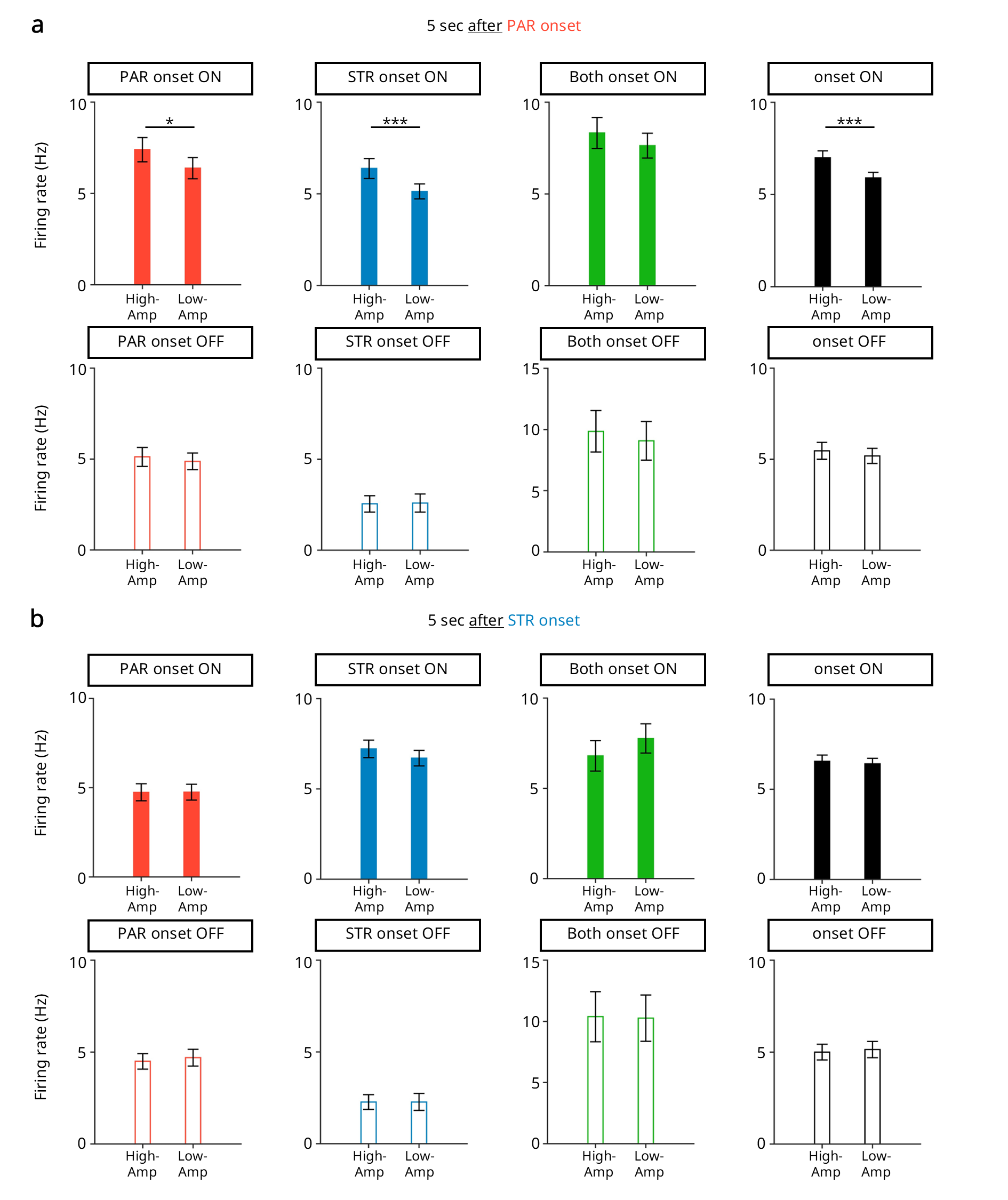

### Figure S5

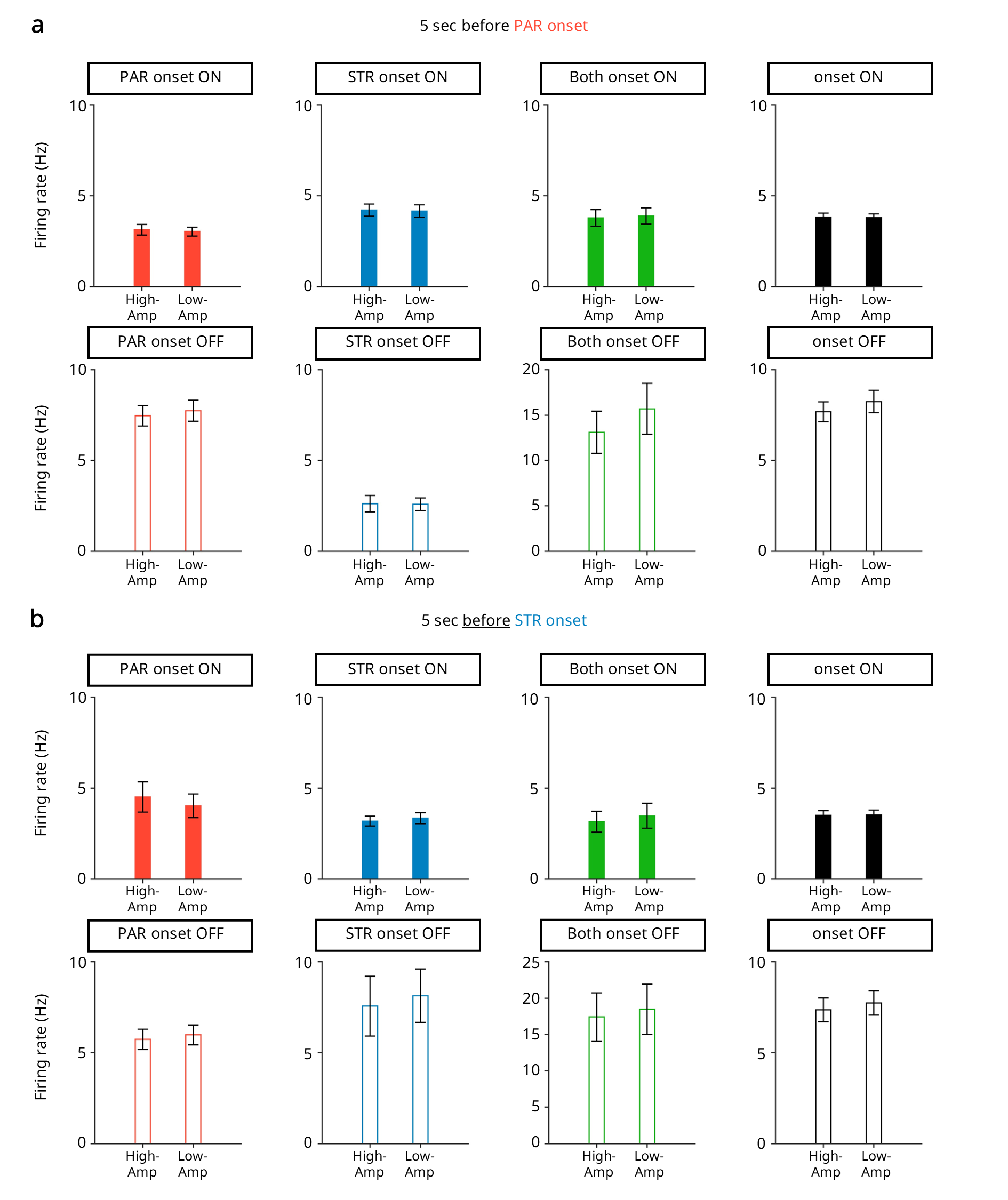
